# Acute cortical stroke alters neural activity in subthalamic nucleus, which correlates with motor disability in rats

**DOI:** 10.1101/2025.01.16.632703

**Authors:** Zhengdao Deng, Myles MC Laughlin, Ugur Kilic, Boateng Asamoah, Bart Nuttin

## Abstract

**Objectives:** We aimed to investigate the impact of acute cortical stroke (ACS) on neural activity in subthalamic nucleus (STN). We then examined the correlation between changes in STN activity and motor disability.

**Methods:** Forty-four Sprague-Dawley rats were used. While rats were anesthetized, we inserted electrodes in STN and induced an ACS by creating photothrombotic lesion in ipsilateral motor cortex. Local field potentials were recorded before and after ACS. The motor behavior was assessed before and after ACS using single pellet reaching task.

**Results:** Rats experienced significant motor disability after ACS. STN firing rate significantly decreased after ACS. Additionally, delta (0.5-4 Hz) and gamma (50-140 Hz) power significantly decreased after ACS. Furthermore, the decrease in delta mean power correlated with decreases in success rate (r =0.77, *p* =0.009) and first try success rate (r =0.69, *p* =0.028). The decreases in gamma mean power (r =0.68, *p* =0.029) and gamma peak power (r =0.74, *p* =0.015) correlated with the decrease in success rate. The decrease in gamma power significantly correlated with the decreased STN firing rate. However, decreased delta power exhibited no correlation with decreased gamma power.

**Interpretation:** ACS causes abnormal STN activity, which correlated with motor disability. Post-stroke STN inhibition may partially compensate for ACS. However, it could also lead to pathological consequences. This STN abnormal activity may serve as a biomarker for motor disability severity after ACS. Furthermore, our findings may provide a possibility for developing neuromodulation strategies, allowing to mitigate post-stroke motor disability through modulating abnormal STN activity.

## Introduction

Stroke is the second leading cause of death, claiming nearly 7 million lives worldwide.^1^ Furthermore, stroke is the third leading cause of motor disability globally.^1^ Approximately 80% of patients with acute stroke experience motor disability in the upper extremities, and 50-60% of these survivors continue to have long-term motor disability.^2^ Hence, it is crucial to gain a deeper understanding of the underlying mechanisms driving post-stroke motor disability. This knowledge is important for further development of effective therapeutic strategies aimed at motor restoration after stroke.

Most therapeutic surgical strategies for post-stroke motor restoration concentrate on targeting motor cortex (MC). The primary approaches aim to normalize cortical abnormalities or strengthen cortical plasticity to reduce motor disability and promote post-stroke motor restoration.^3^ However, their efficacies remain unsatisfactory. Abnormal inter-hemispheric inhibition (IHI) is a possible mechanism underlying motor disability after stroke.^4^ Repetitive transcranial magnetic stimulation (rTMS) was proposed as a therapy to modulate IHI.^4^ However, a Cochrane review indicated that rTMS yields no statistically significant benefits for motor disability.^5^ Neuroplasticity can be enhanced through motor cortex stimulation, which enables motor recovery following stroke.^6^ Levy et al. applied electrical epidural motor cortex stimulation to treat stroke patients. However, their primary results were negative.^7^ Furthermore, anodal transcranial direct current stimulation (tDCS) demonstrated no benefits for motor restoration in stroke patients, despite reports that anodal tDCS enables enhancement of cortical plasticity.^8^ Similarly, while cathodal tDCS inhibits cortical spreading depression, it exerts no positive effects on post-stroke motor restoration.^9^ Despite extensive research on the MC for post-stroke motor restoration, progress in achieving significant motor recovery in humans has been limited. Therefore, it is reasonable to shift some focus from MC to other brain regions that may play a crucial role in post-stroke motor restoration.

The subthalamic nucleus (STN) is a key site for functional convergence of motor circuits and plays a crucial role in motor control and regulation.^10, 11^ It is functionally interlinked with MC through two key pathways: the hyperdirect pathway (HDP) and the STN-cortex pathway (Fig. 1).^12, 13^ These pathways may contribute to motor disability following cortical stroke. Specifically, in abnormal situations such as cortical stroke, abnormal post-stroke intercommunication between the MC and STN through these pathways may lead to motor disability (Fig. 1B), as suggested by Degos’s study.^13^ Cortical delta oscillations are prokinetic and typically decrease after cortical stroke.^14, 15^ This decrease may detrimentally influence or even abnormally synchronize with STN through HDP after cortical stroke. Consequently, STN could become dysfunctional in its role in motor regulation after cortical stroke, leading to motor disability. Additionally, previous studies have shown that STN gamma oscillation is tightly linked to motor function and is prokinetic.^16, 17^ Following cortical stroke, STN gamma oscillation may exhibit abnormal patterns, contributing to motor disability. Moreover, post-stroke abnormal STN oscillations could spread to the MC via STN-cortex pathway, further amplifying the abnormality in MC. This process could exacerbate motor disability and impede post-stroke motor restoration. Ultimately, MC and STN may form an abnormal post-stroke motor circuit after cortical stroke. This circuit may facilitate abnormal post-stroke intercommunication between MC and STN (Fig. 1B), playing a central role in the development and persistence of motor disability after cortical stroke. Therefore, it is crucial to investigate STN neural activity following cortical stroke to better understand its role in post-stroke motor disability. This knowledge may serve as a foundation for developing effective strategies to promote motor restoration after cortical stroke.

**Fig.1.**
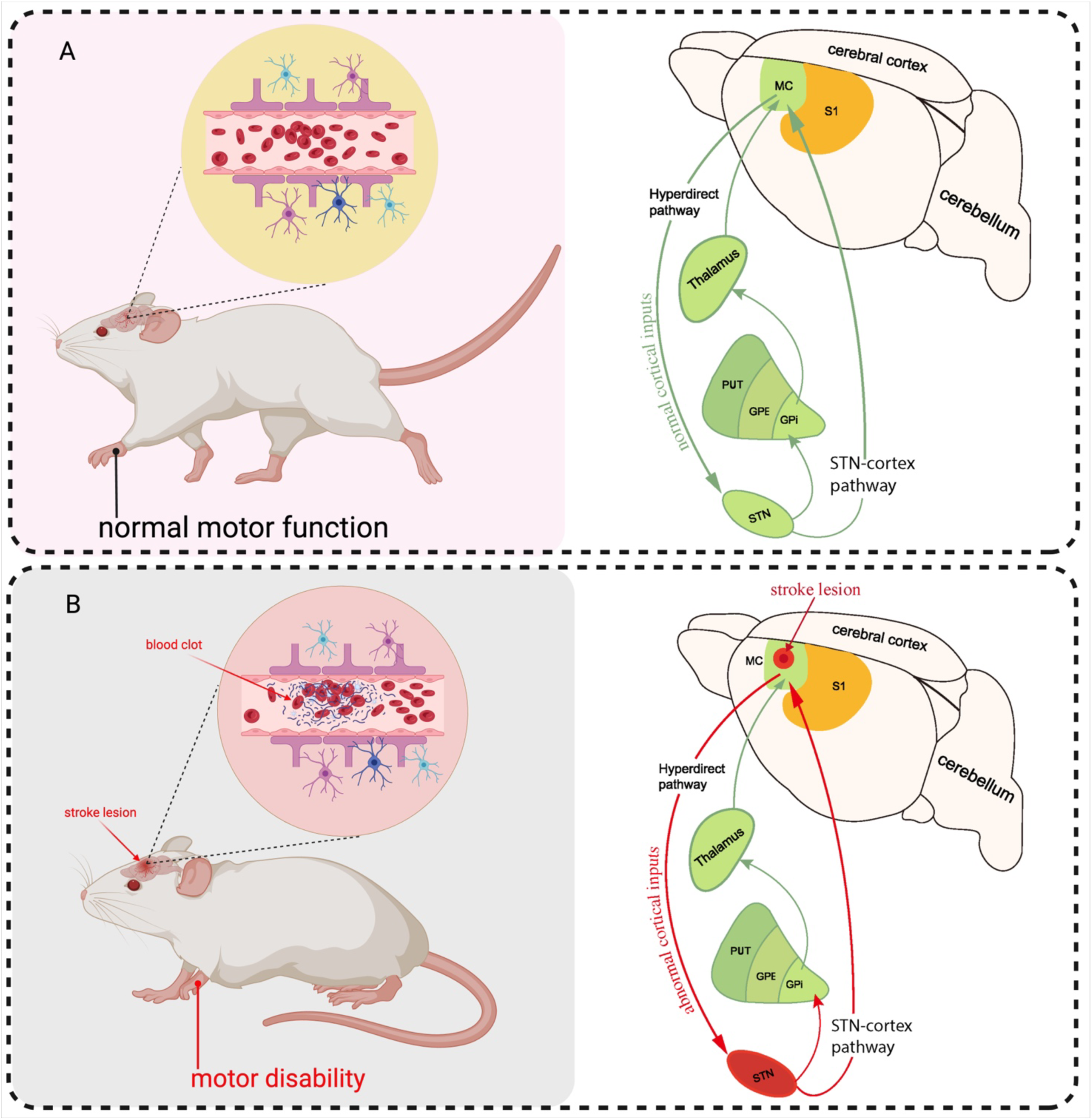
Hyperdirect pathway (HDP) and STN-cortex pathway under normal and acute cortical stroke conditions. (A) The rat exhibits normal motor function with no stroke in MC. MC directly modulates STN and basal ganglion through HDP. The STN projects feedback information to MC via the STN-cortex pathway. Consequently, MC and STN form a functionally integrated network that facilitates interactive communication to regulate motor function. (B) The rat exhibits motor disability after cortical stroke. The cortical stroke causes cortical abnormal activity which propagates to STN via HDP, resulting in an abnormal state in STN. The dysfunctional STN subsequently affects the basal ganglia via the indirect pathway and projects abnormal information to MC via the STN-cortex pathway, thereby exacerbating the abnormal state of the MC. Consequently, the MC and STN may form a post-stroke abnormal motor circuit, facilitating abnormal interactive communication between STN and MC that contributes to motor disabilities following cortical stroke. The green arrows represent normal neural information transmission, while the green brain structures indicate regions that are likely not abnormally modulated by the inputs. In contrast, the red arrows signify abnormal neural information transmission, and the red brain structures indicate regions that may be abnormally modulated by the inputs. MC: motor cortex. HDP: hyperdirect pathway. STN: subthalamic nucleus. S1: primary somatosensory cortex. PUT: putamen. GPE: external globus pallidus. GPi: internal globus pallidus.

In this study, we hypothesised that acute cortical stroke (ACS) causes abnormal neural activity in STN, and this abnormal activity correlates with motor disability. We assessed the motor function using the single pellet reaching task and recorded from STN before and after ACS. Our results revealed that ACS causes significant inhibition of STN neural dynamics in rats. Notably, this inhibition correlates with motor disabilities caused by ACS.

## Materials and Methods

### Animals

All experiments were performed in adult male Sprague-Dawley rats (n=44, 290-340g, Charles River Laboratories). Experiments were carried out in accordance with protocol (P091/2021) approved by KU Leuven Ethical Committee for Animal Experimentation and in accordance with Belgian legislation (Royal Decree regarding the protection of laboratory animals of 29 May 2013) and European directive (2010/63/EU).

### Experimental design

In this study we performed three experiments. Experiment 1: We aimed to determine whether ACS lesion can be reproducibly induced using our modified protocol, based on established method (Fig. 2).^18^ To achieve this goal, we used photothrombotic stroke model. The MC was exposed to a laser beam following an injection of Rose Bengal into tail vein in 10 rats. Next, we studied the variability of the infarct volume ratio of cortical stroke lesions. Experiment 2: We aimed to investigate whether ACS causes alterations in STN firing rate and oscillations, and to determine whether these alterations correlate with motor disability. To achieve this goal, we included an experimental group (n=15) and a sham group (n=15). Before ACS induction, rats in both groups were trained in a single pellet reaching task (SPRT) to assess baseline forelimb motor function (Fig. 3). Baseline STN activity was recorded following electrode implantation in STN. The experimental group underwent a cortical stroke induction, while the sham group received a sham cortical stroke induction (Fig. 4). Post-induction, STN activity was recorded again. In both groups forelimb motor function was re-evaluated 3 days after surgery. This design enabled investigation of ACS-induced changes in STN (Fig. 5) and their potential correlation with motor disability. Experiment 3: We aimed to observe the temporary dynamics of STN activity before, during, and after cortical ACS induction and sham ACS induction across three key phases: pre-induction, during induction, and post-induction of both cortical stroke and sham cortical stroke (Fig. 6). To achieve this goal, four rats were randomly allocated into two groups: an experimental group (n=2) and a sham group (n=2). In both groups, rats underwent electrode implantation into the STN for electrophysiological recording. Rose Bengal was administered at the dosage of 30mg/kg through tail vein. Continuous STN recordings were initiated 10 minutes after the injection and lasted for 16 minutes. The laser remained active throughout these three phases to maintain a consistent recording environment. Specifically, in the experimental group, the laser was operating but the laser beam was fully blocked on its path to MC during the first and last 3 minutes of recording. In the sham group, the laser was also operating but the laser beam was fully blocked on its path to MC during the whole session.

**Fig.2.**
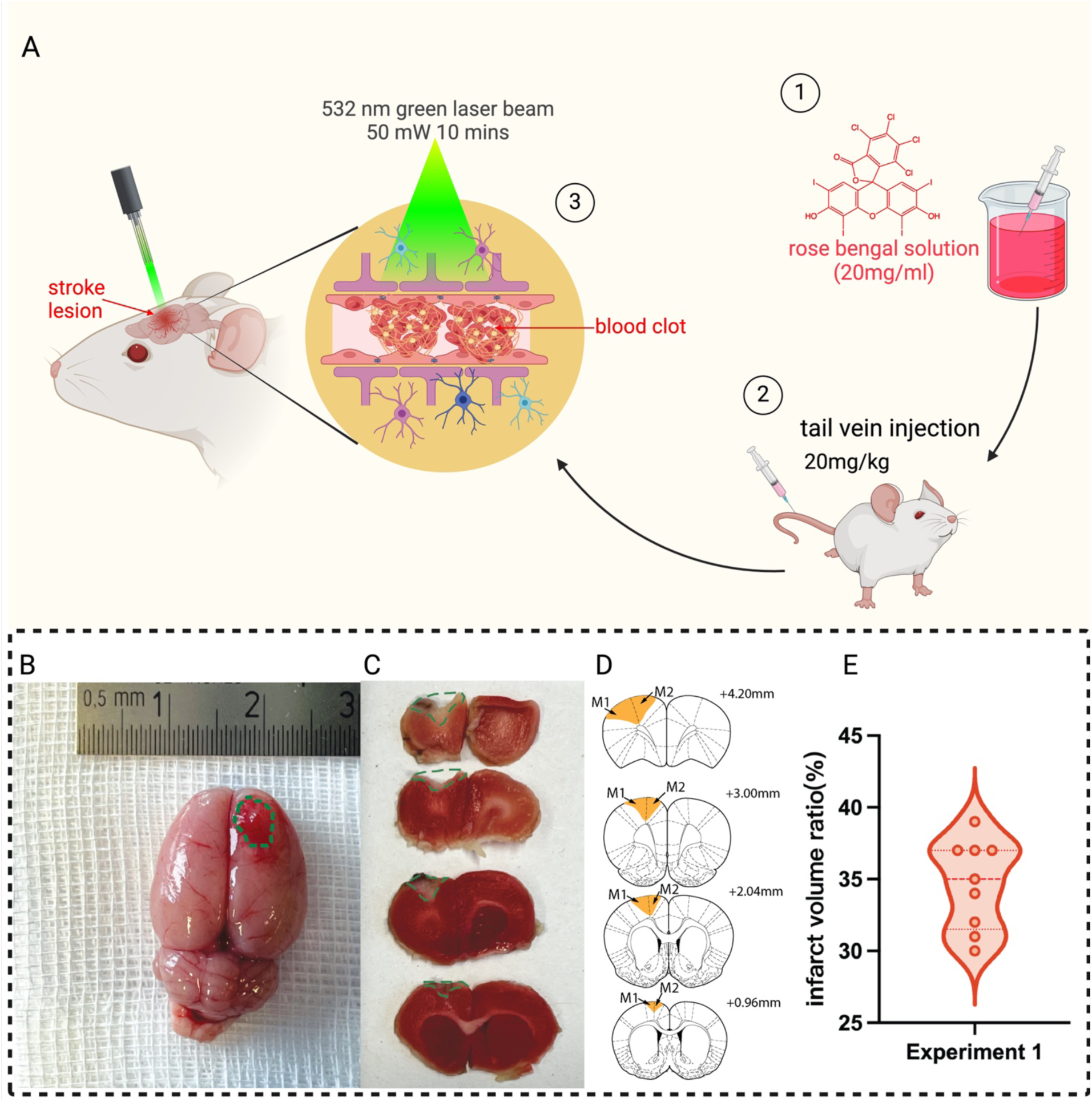
Photothrombotic stroke (ACS) lesion is reliably created in motor cortex (MC) of rats using our protocol. (A) Schematic diagram of creating ACS lesion through our protocol. 1) Rose Bengal is prepared at the concentration of 20mg/ml, 2) and administered to the rats through the tail vein at the dosage of 20 mg/kg. 3) a 4mm-diameter 532nm laser ray (50mW) is directed onto MC for 10 minutes to induce the cortical stroke. (B) Fresh brain tissue collected 24h after cortical stroke induction. The cortical stroke lesion area is circled by green line. (C) 2,3,5-triphenyl tetrazolium chloride (TTC) staining. It shows the cortical stroke lesions induced in each coronal section of a random rat. The cortical stroke lesion is highlighted in green line. (D) Position of the cortical stroke lesion. It shows the position of the cortical stroke lesion in each coronal section shown in Fig. 2C. The lesions, highlighted in yellow, are primarily located in M1 and M2 regions, ranging from 0.96 mm to 4.20 mm anterior to the bregma (Paxinos and Watson’s atlas, 6^th^ edition). M1: primacy motor cortex. M2: secondary motor cortex. (E) Distribution of infarct volume ratio (n=9). This violin plot combines a box plot and a kernel density plot to illustrate the distribution of the infarct volume ratio. The width of the kernel density plot reflects the frequency of data points, with orange dots representing the infarct volume ratio for each individual rat. The mean infarct volume ratio is 34.7 % 3.1%.

### Induction of ACS lesion in MC

General anesthesia was induced by peritoneal injection of a mixture consisting of 0.6 ml ketamine (100 mg/ml, Dechra Pharmaceuticals PLC) and 0.4 ml medetomidine (1 mg/ml, Orion Corporation), administered at a dose of 0.1 ml/100 mg of body weight. The anesthesia was maintained with an injection of the same mixture at a dosage of 0.1 ml/hour. Rats were settled in a stereotaxic apparatus (RWD Life Sciences, China). A midline scalp incision was performed to expose the skull, followed by the creation of a cranial window over right MC for ACS induction. To induce ACS, Rose Bengal (20mg/ml in 0.9% saline, Sigma–Aldrich, Saint Louis, MO, USA) was administered to the rats through the tail vein at the dosage of 20 mg/kg (Fig.2A). Immediately after the injection, a 4mm-diameter 532nm laser ray (50mW, Shanghai MRTRADE International Co., LTD, China) was directed onto MC for 10 minutes. After cortical stroke induction, rats were removed from the stereotaxic apparatus and returned to the cage on a heating pad for recovery. Then meloxicam (2mg/kg; Metacam, Boehringer Ingelheim, Germany) was administered subcutaneously once for post-operative pain care.

### Measurement of the ACS lesions

In experiment 1, after 24 hours of ACS induction, rats were euthanized, and fresh brain tissues (Fig.2B) were collected and frozen at −18 °C for 25 minutes. Following the freezing process, the brain tissue was cut into coronal sections (1 mm) using a rat brain matrix (Agnthos, Sweden). The sections containing infarct lesion were incubated in 2% 2,3,5-triphenyl tetrazolium chloride solution (TTC, Sigma-Aldrich, USA) at 37 °C for 30 minutes. After incubation, the sections were fixed in 4% paraformaldehyde for 24 hours and photographed (GoPro Hero 11, USA). The infarct region was identified as the stroke cavity together with the surrounding white area stained by TTC in MC (Fig.2).^18^ The normal brain tissue was identified as the rest of the brain (i.e. excluding the infarct region). The TTC-stained sections were further analyzed using ImageJ (NIH Image, USA). We expressed the infarct volume ratio (1) as in Mukda et al.^19^ by calculating the ratio of infarct volume to normal brain tissue volume. Results were given as the mean ± standard deviation of the infarct volume ratio.

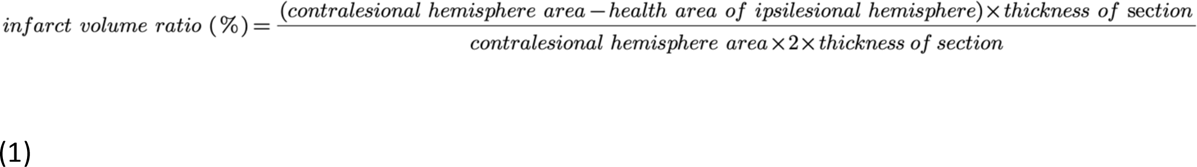

### SPRT and motor evaluation

SPRT was conducted based on Whishaw’ et al.^20^ Rats were trained in a Plexiglas chamber measuring 15cm (width) × 24cm (length) × 36cm (height), featuring a narrow slit (10 mm wide, 15 mm high) at the center of the front wall of the chamber. An exterior shelf, standing 3 cm high, was attached to the front wall to hold chocolate pellets (45mg, Bioserv Corporation, USA). The pellet was positioned in an indentation 15 mm from the chamber’s outer wall (Fig. 3B, C), ensuring that the rat could not retrieve it with its tongue.

**Fig.3.**
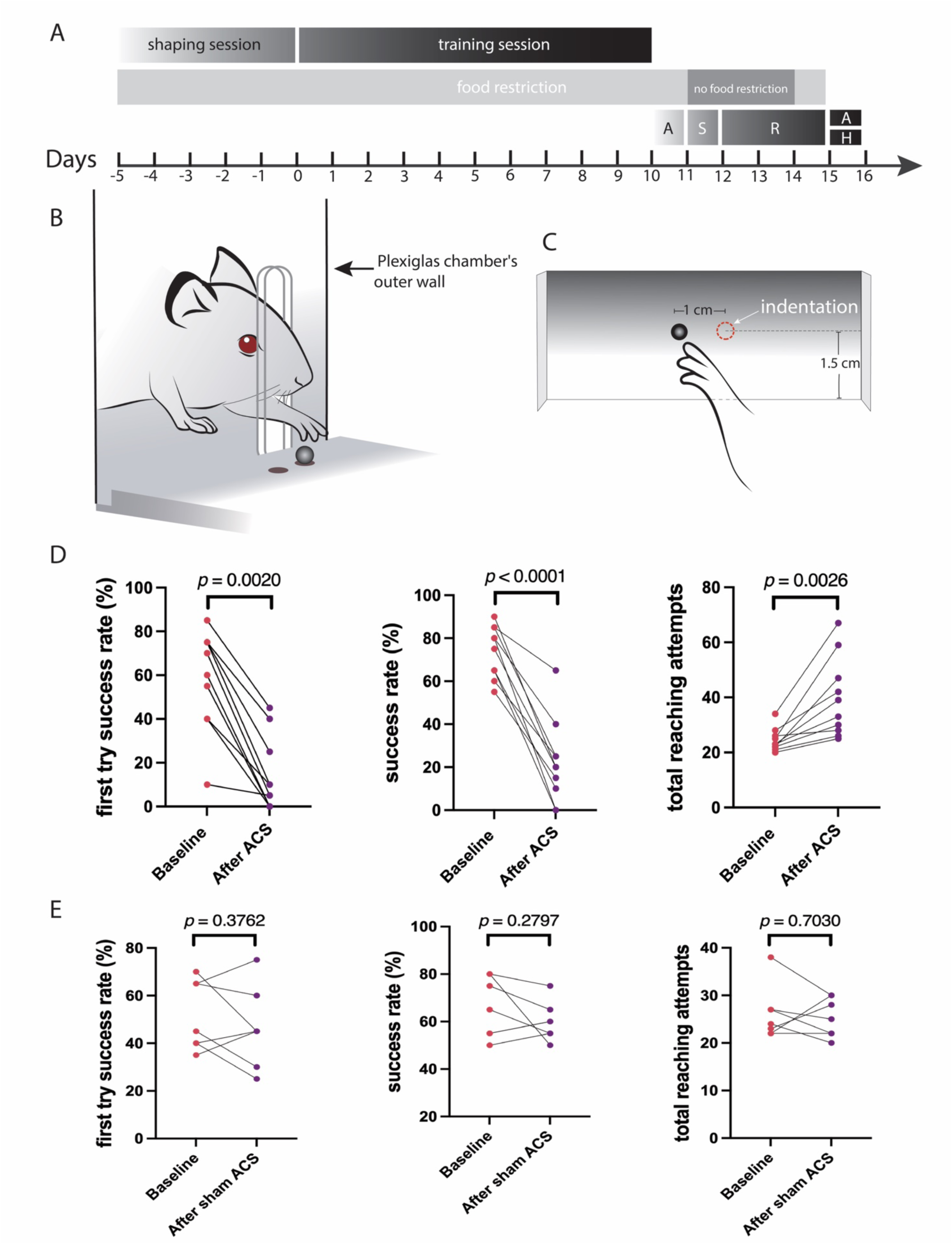
Single pellet reaching task (SPRT) and motor evaluation. (A) Experimental design for Experiment 2. The flowchart shows that rats underwent SPRT training consisting of 5-day shaping and 10-day training session. After completing the training, the rats were subjected to motor evaluations, surgical procedures, and subsequent histological analysis. A: assessment of motor function. S: surgery. R: recovery. H: histological analysis. (B) Schematic diagram illustrating the apparatus for SPRT. Rat is being trained in a Plexiglas chamber featuring a narrow opening, allowing it to reach through and retrieve food pellets. (C) A detailed close-up of the food pellet shelf of the apparatus for SPRT. The food pellet is positioned in one of the indentations 1.5 cm from the Plexiglas chamber’s outer wall. The distance between the indentations center point is 1cm. (D) First try success rate and success rate significantly decreased after acute cortical stroke (ACS) induction (n=10). The success rate decreased from 74% to 23% (*p* < 0.0001, Student’s *t*-test), the first try success rate decreased from 59% to 14% (*p* = 0.0020, Wilcoxon signed-rank test) after ACS. The total reaching attempts significantly increased from 25 to 40 (*p* = 0.0026, Student’s *t*-test) after ACS. (E) First try success rate, success rate and total reaching attempts remained unchanged after sham ACS induction (n=7). No significant differences in first try success rate (*p=* 0.3762), success rates (*p=* 0.2797) or total reaching attempts (*p=* 0.7030) before and after sham ACS induction. Statistics was applied using Student’s *t*-test.

SPRT training consisted of a 5-day shaping session (SS) followed by a 10-day training session (TS) (Fig. 3A). The rats were food restricted before executing SPRT training, and their body weight remained at approximately 90% of their original weight during SPRT session. During SS, 50 food pellets were provided daily on the rat cage floor to help rats acclimate to consuming chocolate pellets. On SS Day 1-2, rats were introduced to the Plexiglas chamber for 30 minutes to acclimate to the environment. Starting from SS Day 3, chocolate pellets were positioned in front of the central slit, allowing the rats to retrieve the chocolate pellets on the shelf. The pellets were gradually placed farther away on the shelf until the rats extended their forelimbs to reach through the slit and retrieve the chocolate pellets in the indentations. On SS Day 5, the chocolate pellets were positioned in indentations, allowing rats to retrieve them and demonstrate forelimb dominance (Fig. 3C). The dominant forelimb is defined as the one the rats prefer to use at least 70% of the reaches. During TS, rats were conditioned to walk from the back to the front of the chamber to retrieve a chocolate pellet in the indentation. They completed 60 trials in each daily session. Rats were trained 10 days to retrieve chocolate pellets with their dominant forelimb. Forelimb motor function was evaluated 1 day before stroke induction (baseline) and 3 days after. A camera (GoPro Hero 11, USA) was used while rats were conducting SPRT for motor assessment. Rats performed 20 trials during motor evaluation. A successful trial was counted when a rat grasped a chocolate pellet and consumed it. If the rat succeeded on the first attempt, it was counted as a first successful trial. We used three endpoint measurements: success rate (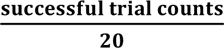 × 100%), first-attempt success (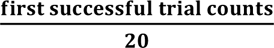 × 100%), and total number of reaching attempts.^21^ The rats that achieved at least 50% success rate were used in further experiments.

### Electrode implantation into STN

A midline scalp incision was performed to expose the skull. Micro-electrodes (Õ = 125 μm, FHC, USA) composed of platinum/iridium (200–300 kΩ) were used for recording. In experiment 2, we implanted the electrode in STN contralateral to the dominant forelimb. In experiment 3, we implanted the electrode in the right STN. The STN coordinates were taken as the average of bregma and interaural coordinates: 3.7mm posterior, 2.4mm lateral and 8.35mm ventral to bregma. 5.3mm anterior, 2.4mm lateral and 1.6mm dorsal to interaural point.^22, 23^ Dental cement was used to affix the electrode to the skull. Eight stainless steel bone screws (1.19 mm, Fine Science Tools, USA) were inserted into boreholes, while ensuring the dura mater remained intact, to anchor the dental cement to the skull in the subsequent step. A wire connected to a screw in the occipital bone served as the reference electrode during electrophysiological recordings. Additional dental cement was applied to the bone screws to form a cap to protect the electrode. Following electrode implantation and electrophysiological recordings, the wound edges were closed with sutures at both the front and back of the cap, and the rats were returned to their cages.

### STN recordings

In Experiment 2, the STN contralateral to the dominant forelimb was recorded. Recordings were conducted before and after ACS in the experimental group and sham group. Each recording session lasted 3 minutes. The baseline recording in both groups was taken 15 minutes after electrode implantation in STN. (Fig.4A and 4E). This was completed before creating a cranial window over the MC region to prevent potential irritation or micro-lesions to MC during skull drilling in both groups, which might potentially affect STN oscillations and contaminate the baseline recording. ACS and sham ACS were induced in MC contralateral to the dominant forelimb after Rose Bengal injection. This was completed according to the protocol described previously in “Induction of ACS lesion in MC” for both the experimental and sham groups. For sham ACS stroke induction, the laser was operating, while its laser beam was entirely blocked by an impenetrable tape (Tesa, Germany) placed over the laser outlet to prevent it from reaching MC. STN recording was performed again 30 minutes after stroke induction in experimental group (Fig.4C) and sham group (Fig.4E). In Experiment 3, ACS was induced in right MC in experimental group, while a sham ACS was induced in control group. Continuous right STN recording was performed for both the experimental and sham groups. Recording began 10 minutes after the injection of Rose Bengal and lasted for 16 minutes in both groups. The laser was operating throughout the entire recording period, but during the first and last three minutes of recording, the laser beam was entirely blocked in the experimental group. In the sham group, the laser was also operating throughout the entire recording period, but the beam remained entirely blocked throughout the whole recording period.

**Fig.4.**
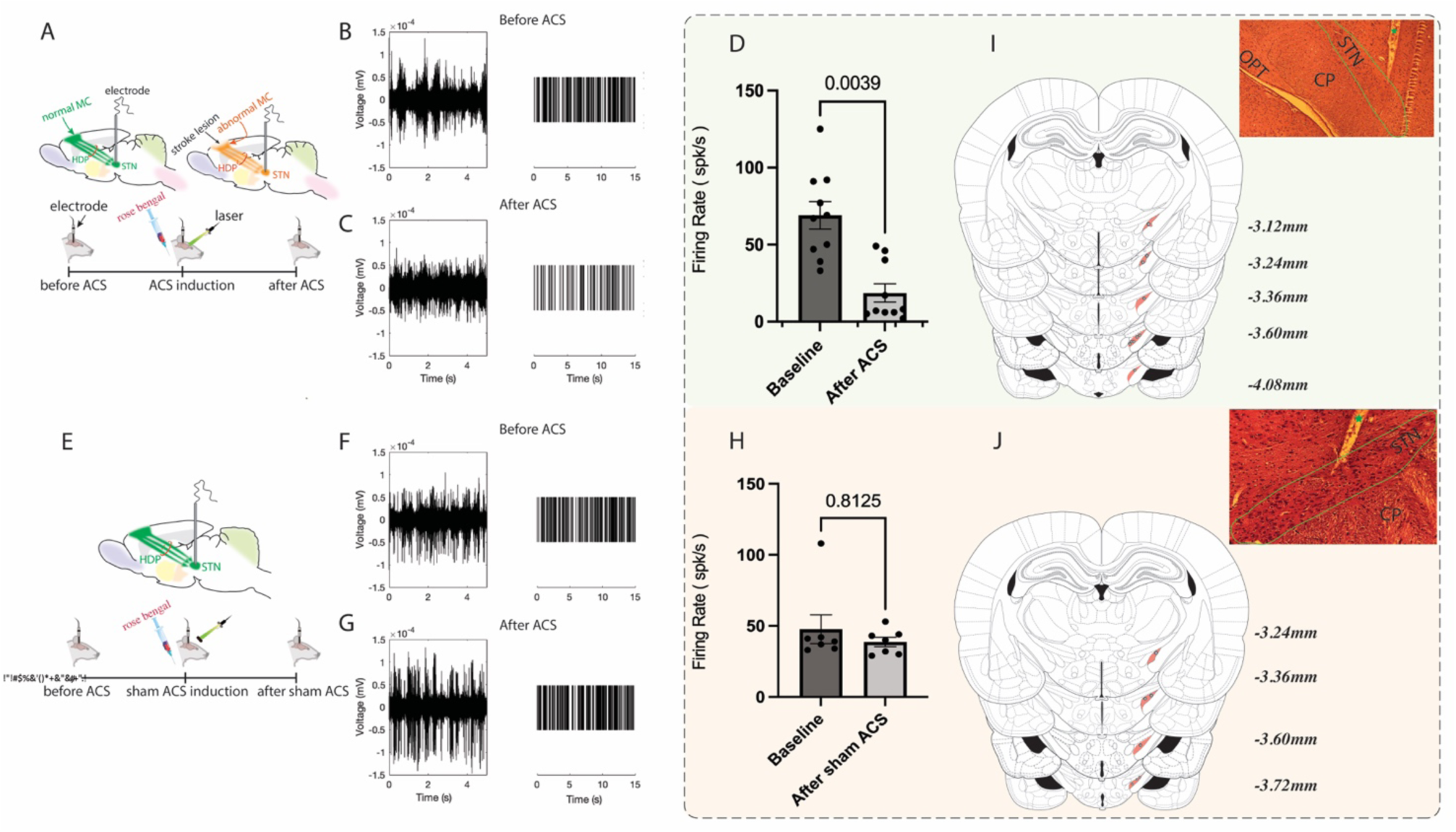
STN firing pattern and firing rate. (A) STN is recorded before and after acute cortical stroke (ACS) induction. (B) The STN firing is characterized by an irregular fast burst-firing pattern, marked by rapid, irregular bursts of high-amplitude spikes, which was consistent with previous studies.^25, 26^ (C) In contrast, the STN firing pattern is noticeably affected by ACS, shifting to a relatively low-frequency burst-firing pattern characterized by irregular bursts of low-amplitude spikes. (D) The results reveal that the STN firing rate significantly decreased from 69 %28 spks/s to 19 % 18 spks/s after ACS induction. Wilcoxon signed rank test (*p*= 0.0039, Wilcoxon signed-rank test, n=10). (E) STN is recorded before and after sham ACS induction in control group. (F,G) The firing pattern remains consistent before and after the sham ACS induction. (H) The results reveal that the STN firing rate remained unchanged after sham ACS induction (*p=*0.8125, Wilcoxon signed-rank test, n=7). The position of each electrode tip in experimental group (I) and control group (J) of Experiment 2 are showed in the reconstructed schematics based on the Paxinos and Watson’s atlas (6^th^ edition). The position of each electrode tip is in STN, ranging from 3.12 mm to 4.08 mm posterior to the bregma in experimental group, and 3.24 mm to 3.72 mm posterior to the bregma in control group. The image shows the position of electrode tip in STN of a random rat. STN is circled by the green line. The electrode trajectory is highlighted by a green asterisk. HDP: hyperdirect pathway. MC: motor cortex. STN: subthalamic nucleus. OPT: optic tract. CP: cerebral peduncle.

### Electrophysiological data acquisition

The signals recorded from the micro-electrodes were amplified, bandpass filtered (0.1 Hz–7.9 kHz), and digitized (16-bit, 30 kHz). This process was facilitated by an RHD 32 Intan head-stage (Intan Technologies, Los Angeles, CA) and an Open Ephys Acquisition Board (open-ephys.org). Neural activity was visualised through Open Ephys GUI.

### Histological verification of the electrode tip

At the end of experiments, rats were anaesthetized and underwent cardiac perfusion. The brain sample was quickly extracted and immersed in 4% paraformaldehyde for paraffin embedding. Subsequently, the brain was sectioned into 10 μm slices and stained using hematoxylin and eosin (H&E). The position of electrode tip was verified using H&E staining.

### Power Spectral Density (PSD)

In experiment 2, we aimed to investigate whether STN oscillations in the delta (0.5-4 Hz) and gamma (50-140 Hz) frequency bands changed after ACS. To address this, we performed spectral analysis to compute the PSD in delta (0.5-4 Hz) and gamma (50-140 Hz) bands. Raw data were collected at a sampling rate of 30 kHz and visually inspected to ensure signal quality prior to analysis. Spectral analysis was conducted using short-time Fourier transforms, implemented through MATLAB’s spectrogram function, with a 5-second hanning window, 50% overlap, and a 5-second FFT length. We quantified the mean power, peak power, and peak frequency in the oscillation bands both before and after ACS induction. Mean power was defined as the average PSD across the bands, calculated using the formula: 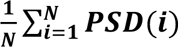, where *N* is the number of 5 second time windows, and *PSD(i)* represents the PSD value at each frequency bin. Peak power was defined as the maximum PSD value in each band, and peak frequency was the frequency corresponding to this peak power. To effectively visualise the difference of gamma band before and after ACS, the PSD values in this band were converted to a logarithmic scale by using the formula 10 · *log*_10_(*PSD*), and smoothed using a moving average filter. In experiment 3, we aimed to calculate the PSD in the delta (0.5-4 Hz) and gamma (50-140 Hz) frequency bands over time. To do this, the spectral analysis was conducted.

For each recording session, we calculated the mean delta and gamma PSD at each time point using the formula: 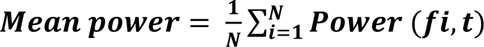, where power (*fi, t*) is the PSD at frequency *fi* and time t, N is the number of frequency bins in the frequency band. To effectively visualize delta and gamma bands, the PSD values were converted to a logarithmic scale and smoothed. We aggregated the session data into a pool, analyzed the subsequent recording session using the same method, and then added the new session data to the pool for further process. Finally, the *mean power* was averaged across all sessions (n=2).

### Statistical Analysis

In experiment 1, we aimed to analyze the variability of the ACS lesions created using our protocol. Thus, we used coefficient of variation (CV), calculated as the ratio of the standard deviation of infarct size to the mean infarct size to represent the variability of the ACS lesions. A CV of less than 10% indicates a high reproducibility.^23^

In experiment 2, we aimed to evaluate whether motor function measures changed before and after ACS. To achieve this, we first assessed the normality of the datasets using the Kolmogorov-Smirnov test. A *p*-value < 0.05 indicated a deviation from normality, in which case we applied the Wilcoxon signed-rank test. Conversely, a *p*-value > 0.05 suggested normal distribution, in which case we applied the Student’s *t*-test for analysis. Next, we evaluate whether STN firing rate, STN delta (0.5-4 Hz) and gamma (50-140 Hz) oscillations, including peak power, mean power, and peak frequency, changed before and after ACS induction. The appropriate statistical test was selected for each comparison based on the dataset normality. Finally, we aimed to evaluate the correlation between motor disability and the changes in delta (0.5-4 Hz) oscillation on the one hand and gamma (50-140 Hz) oscillations on the other. To achieve this, we first assessed the normality of the datasets using the Kolmogorov-Smirnov test. Since the *p*-value > 0.05, indicating normal distribution, Pearson’s correlation coefficient and simple linear regression were applied for the analysis. Additionally, we aimed to evaluate the correlation between the changes in delta power and the changes in gamma power, the correlation between the decreased STN firing rate and the changes in delta power, the correlation between the decreased STN firing rate and the changes in gamma power. To address this, we utilize Pearson’s correlation coefficient and simple linear regression, as the data were normally distributed, as confirmed by the Kolmogorov-Smirnov test. These analyses were performed using Prism version 9 (GraphPad Software, La Jolla, CA, USA) and MATLAB (R2021b, The MathWorks, Inc., Natick, Massachusetts, USA). All results are presented as mean ± standard deviation. A significance level of *p* < 0.05 was established for statistical significance.

## Results

### ACS lesion was highly reproducible in rats

To enable the application of our protocol for creating ACS lesion in MC for subsequent experiments, we initially validated its reproducibility in Experiment 1, which involved nine rats for further analysis. The results showed that the ACS lesion was primarily localized to the primary and secondary MC (Fig.2C,D). TTC staining revealed that the mean infarct volume ratio was 34.7 ± 3.1% (Fig.2E), with a CV of 8.9%, indicating that ACS was reproducibly induced in MC using our protocol.^24^

### ACS caused a significant inhibition in STN firing rate

ACS lesions were shown to be highly reproducible in MC using our protocol in Experiment 1. Therefore, we employed this approach in all experiments. We initially studied the impacts of ACS on STN activity. STN was recorded before and after ACS induction and sham ACS induction in experimental group and sham group. For the analysis, 10 rats were included in the experimental group and 7 rats in the control group, as these rats successfully underwent electrode implantation surgery. The results showed that the typical STN firing pattern was identified as an irregular fast burst-firing pattern, marked by rapid, irregular bursts of high-amplitude spikes (Fig.4B).^25, 26^ Following ACS induction, however, this firing pattern shifted to a relatively low-frequency burst-firing pattern, marked by irregular bursts of low-amplitude spikes (Fig. 4C). Furthermore, our results revealed that the STN firing rate significantly decreased from 69 ± 28 spks/s to 19 ± 18 spks/s (*p=*0.0039, Wilcoxon signed-rank test, Fig.4D). However, the firing pattern and firing rate (*p =*0.8125, Wilcoxon signed-rank test) did not show changes after sham ACS induction (Fig. 4F, G, H).

### ACS caused a significant decrease of delta and gamma power in STN

STN firing rate was affected by ACS, which raised the question of whether its neural oscillations are also affected by ACS. Since delta and gamma oscillations were reported to be associated with movement, ^12–14^ we further investigated the effects of ACS on STN delta (0.5-4 Hz) and gamma (50-140 Hz) oscillations. After ACS, both STN delta mean power (*p* = 0.0009, Student’s *t*-test) and delta peak power (*p* = 0.0001, Student’s *t*-test) significantly decreased, while the delta peak frequency remained unchanged (*p* = 0.1934, Student’s *t*-test, Fig.5A, E). Similarly, STN gamma mean power (*p* = 0.0004, Student’s *t*-test) and gamma peak power (*p* = 0.0008, Student’s *t*-test) also showed significant decreases, while the peak frequency remained unchanged after ACS (*p* = 0.0840, Wilcoxon signed-rank test, Fig.5B, F). In contrast, no significant changes were observed in the STN delta mean power (*p*=0.2496, Student’s *t*-test), delta peak power (*p*=0.0832, Student’s *t*-test), gamma mean power (*p*>0.99, Wilcoxon signed-rank test), or gamma peak power (*p*=0.8438, Wilcoxon signed-rank test) following sham ACS induction (Fig. 5C, D, G, H).

**Fig.5.**
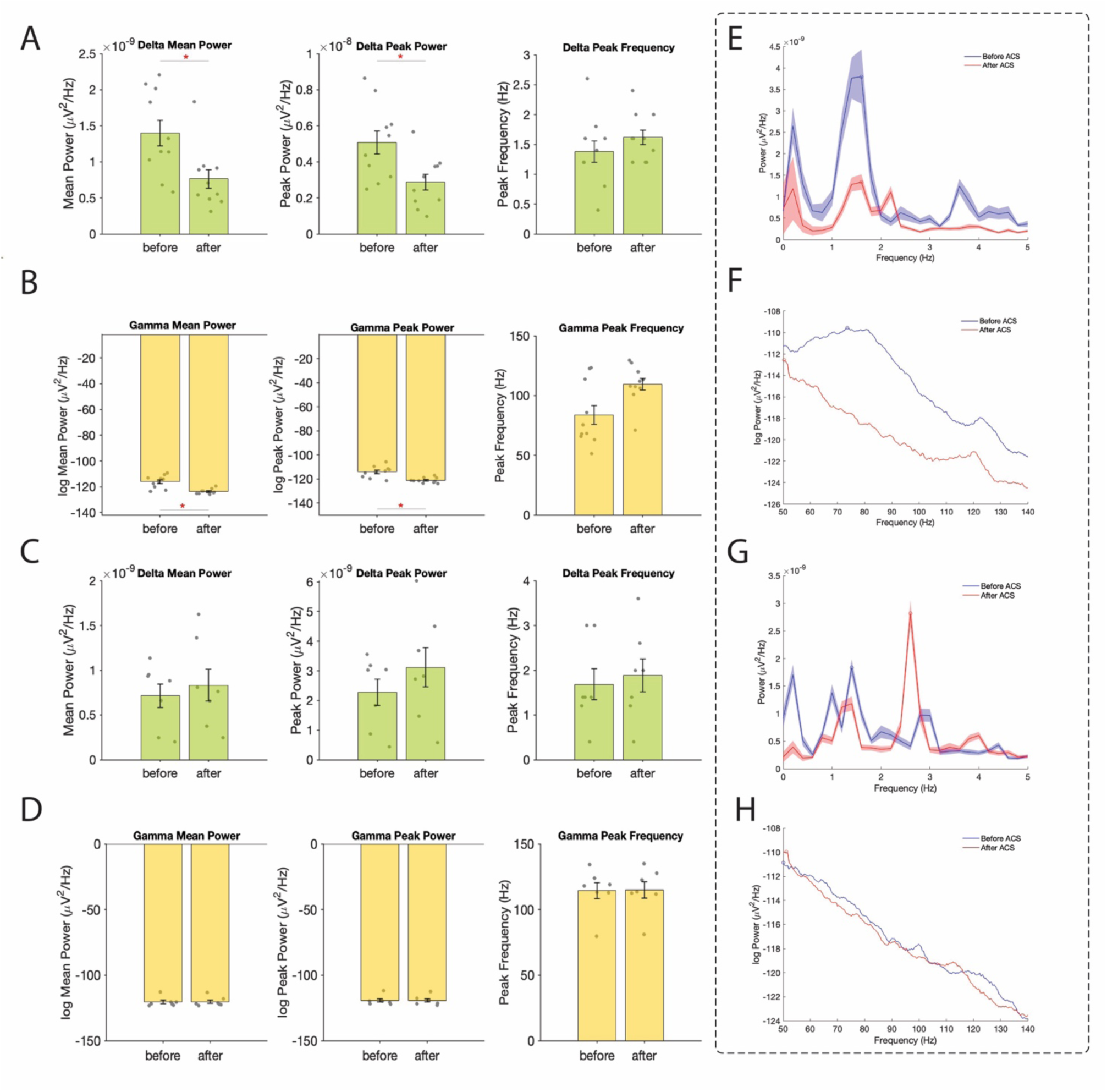
STN delta and gamma oscillations. (A) After acute cortical stroke (ACS) induction, the delta mean power (*p* = 0.0009, Student’s *t*-test) and peak power (*p* = 0.0001, Student’s *t*-test) significantly decrease. However, the delta peak frequency (*p* = 0.1934, Student’s *t*-test) remains unchanged, n=10. *indicates the statistical significance. (B) After ACS induction, the gamma mean (*p* = 0.0004, Student’s *t*-test) and peak power (*p* = 0.0004, Student’s *t*-test) significantly decrease. However, the gamma peak frequency (*p* = 0.0840, Wilcoxon signed-rank test) remains unchanged, n=10. *indicates the statistical significance. (C) After sham ACS induction, the delta mean (*p*=0.2496, Student’s *t*-test) and peak power (*p*=0.0832, Student’s *t*-test), and the delta peak frequency remain unchanged, n=7. (D) After sham ACS induction, the gamma mean power (*p*>0.99, Wilcoxon signed-rank test) and peak power (*p*=0.8125, Wilcoxon signed-rank test) remain unchanged, n=7. (E,F) STN delta (0.5-4 Hz) and gamma (50-140 Hz) oscillations in a random rat in the experimental group of Experiment 2. The delta mean and peak power show a noticeable decrease after ACS induction (E). Similarly, the gamma mean power and peak power decrease after ACS induction (F). (G,H) STN delta (0.5-4 Hz) and gamma (50-140 Hz) oscillations in a random rat in the control group of Experiment 2. It shows the changes of delta power (G) and gamma power (H) before and after ACS induction.

### ACS caused significant forelimb motor disability in rats

STN activity was found to be altered by ACS, indicating disruptions in motor behaviour, given the STN’s critical roles in motor regulation and control.^10, 11^ Therefore, we further tested the rats’ forelimb motor function after ACS. Using the SPRT, we assessed forelimb motor performance both before and after ACS. The results revealed that forelimb motor function was significantly impaired after ACS (Fig.3D). Specifically, the success rate decreased from 74% to 23% (*p* < 0.0001, Student’s *t*-test). Additionally, the first try success rate decreased from 59% to 14% (*p* = 0.0020, Wilcoxon signed-rank test). Conversely, the total reaching attempts significantly increased from 25 to 40 (*p* = 0.0026, Student’s *t*-test) after ACS. In contrast, sham cortical stroke showed no difference in forelimb motor function in rats (Fig. 3E), with no significant differences in success rates (*p=* 0.2797, Student’s *t*-test), first try success rate (*p=* 0.3762, Student’s *t*-test) or total reaching attempts (*p=* 0.7030, Student’s *t*-test) before and after sham ACS induction.

### STN firing rate and oscillation power decreased during ACS induction

STN activity was altered by ACS, raising a question of whether ACS causes this change during the induction process. To answer this, we continuously recorded STN activity throughout the entire ACS induction session in both the experimental and control groups. The results revealed that STN firing rate, delta, and gamma power showed a gradual decrease during and after ACS induction (Fig.6B, C). Notably, STN firing exhibited a fast response to ACS induction, with a noticeable decrease occurring around 20 seconds after ACS induction (n=2). However, STN delta and gamma power exhibited a slow response to ACS induction, with a decrease occurring at approximately 600 seconds after stroke induction (Fig. 6B). In contrast, the STN firing rate, delta, and gamma power did not decrease during or after sham ACS induction (n=2, Fig. 6E, F).

**Fig.6.**
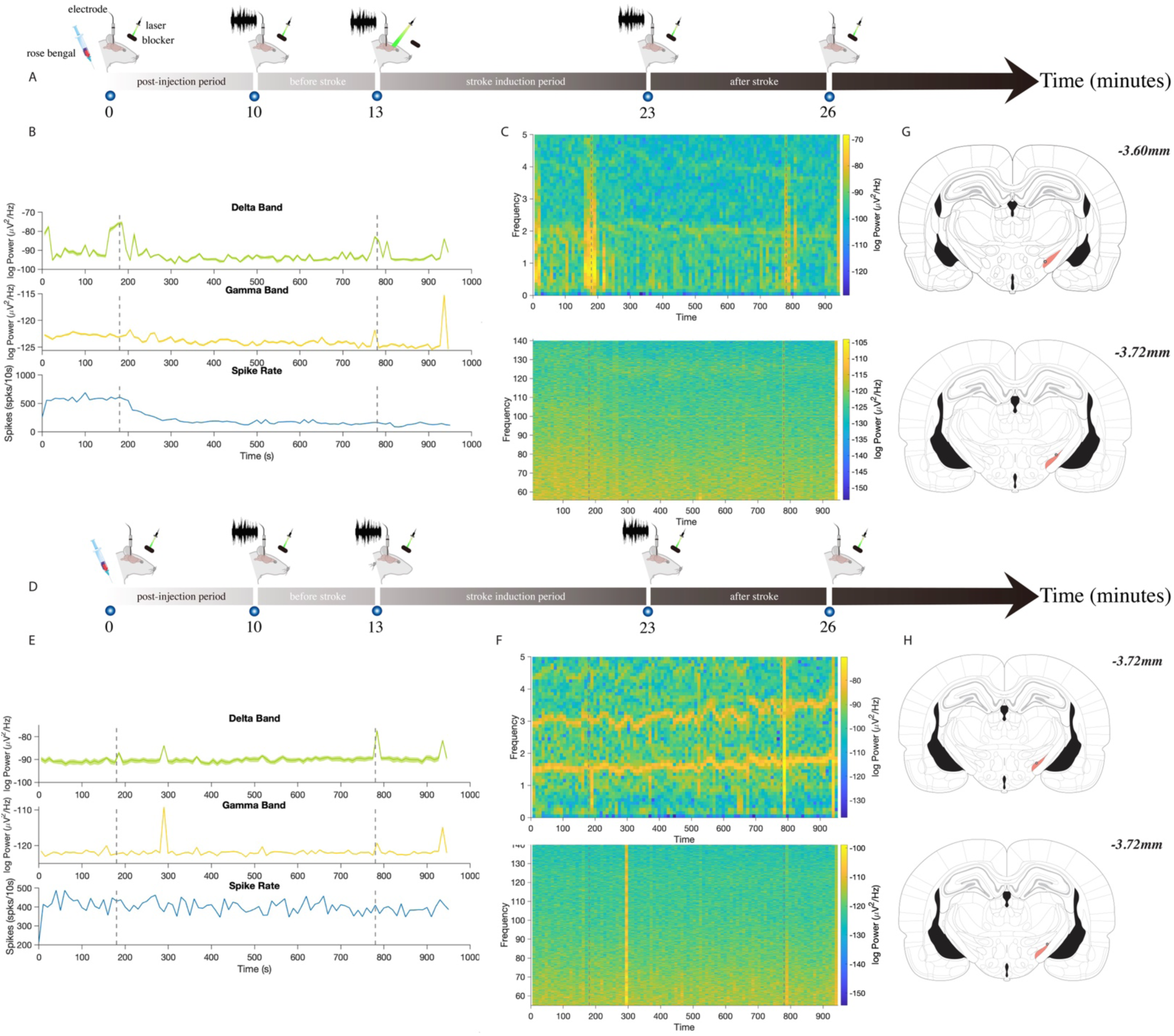
STN dynamics before, during and after stroke and sham stroke induction. (A) Procedures executed in experimental group of Experiment 3. Continuous STN recording was initiated 10 minutes after injection and lasted for 16 minutes. The laser was operating but the laser ray was fully blocked on its path to motor cortex (MC) during the first and last 3 minutes of recording. (B) Dynamics of STN oscillations and firing rate in rats (n=2) before, during and after stroke induction. STN firing responded rapidly to acute cortical stroke (ACS) induction, decreasing around 20 seconds after. In contrast, STN delta and gamma power showed a slower response, decreasing around 600 seconds after induction. (C) Time-frequency analysis. It shows the changes of STN delta (0.5-4 Hz) and gamma (50-140 Hz) oscillations before, during and after ACS induction. (D) Procedures executed in control group of Experiment 3. Continuous STN recording was initiated 10 minutes after injection and lasted for 16 minutes. The laser was operating but the laser ray was fully blocked on its path to MC in the whole session. (E) Dynamics of STN oscillations and firing rate in rats (n=2) before, during and after sham ACS induction. Normal cortical inputs were expected to be continuously captured by the electrode. STN oscillations and firing rate remained stable before, during and after sham ACS induction. (F) Time-frequency analysis. It shows the dynamics of STN delta (0.5-4 Hz) and gamma (50-140Hz) oscillation before, during and after sham ACS induction. (G,H) Histology results. The positions of the electrode tips in rats subjected to ACS (G) and sham ACS (H) induction are shown. STN is highlighted in red area, while the black dots represent the positions of electrode tips. The first and second vertical dotted lines in (B,C,E,F) represent the start and end of the electrophysiological recording.

### STN delta and gamma power positively correlated with motor disability

ACS caused motor disability and a decrease in STN delta and gamma oscillations in the same rats, indicating motor disability might be correlated with these decreased oscillations.^14–16^ Thus, we further analyzed the correlation between the changes in SPRT success rate and first try success rate with the changes in delta and gamma peak power and mean power using Pearson’s correlation test. The results revealed a significant passive correlation between the decrease in STN delta mean power and both the decreased SPRT success rate (r = 0.77, *p* = 0.009) and the first-try success rate (r = 0.69, *p* = 0.028). In contrast, decreased delta peak power did not significantly correlate with both the decreased success rate (r =0.21, *p* =0.565) and first try success rate (r =0.32, *p* =0.376) (Fig. 7A). The Pearson’s correlation test also revealed decreased gamma mean power (r = 0.68, *p* = 0.029) and peak power (r = 0.74, *p* = 0.015) were both positively correlated with decreased success rate (Fig. 7B). However, neither decreased gamma mean power (r = 0.39, *p* = 0.259) nor gamma peak power (r = 0.41, *p* = 0.242) were found to be correlated with first attempt success rate.

**Fig.7.**
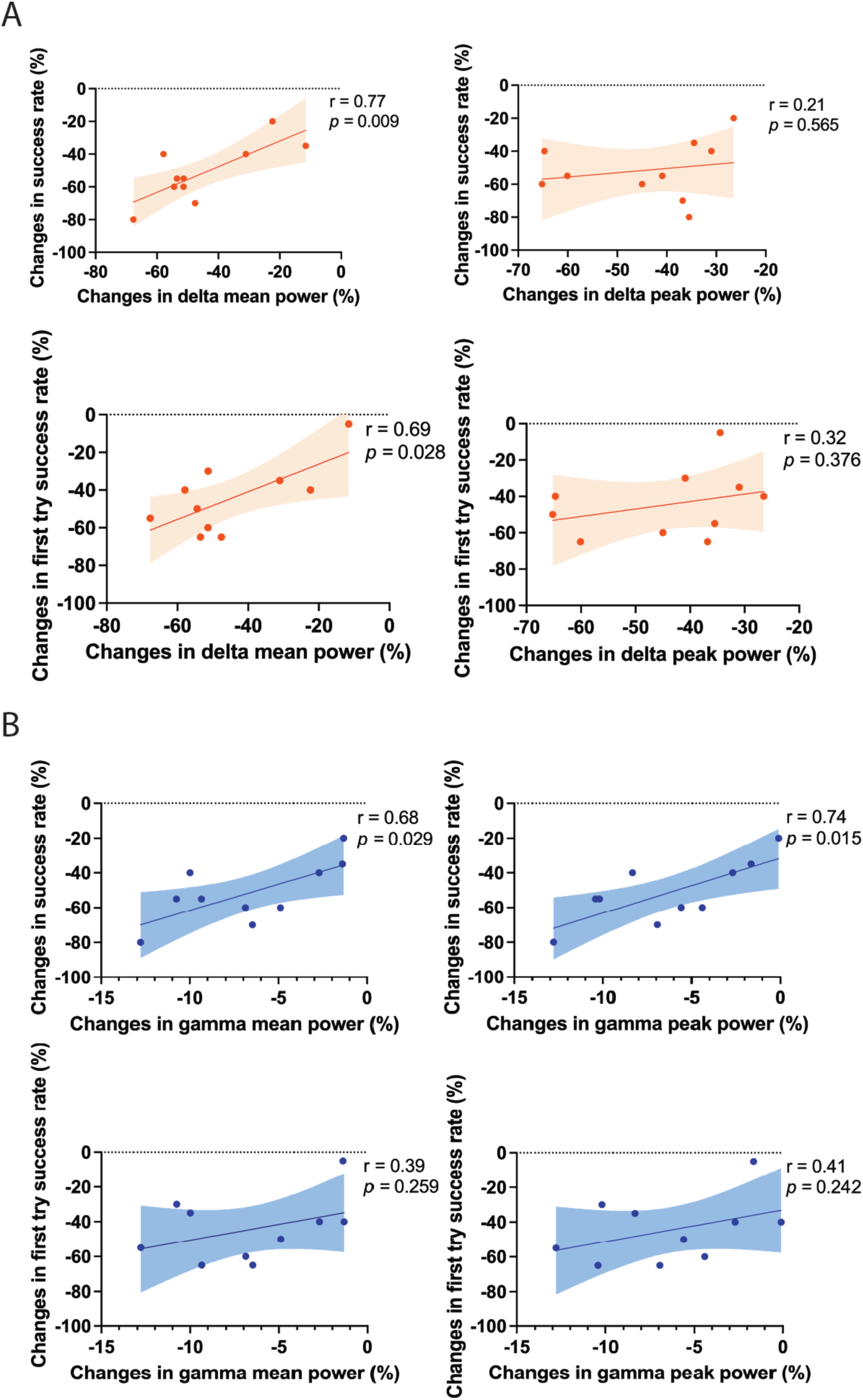
Spearman’s rank correlation results. A. Correlations between changes in STN delta oscillation and motor disability. The results revealed a significant passive correlation between the decrease in STN delta mean power and both the decreased SPRT success rate (r = 0.77, *p* = 0.009) and the first-try success rate (r = 0.69, *p* = 0.028). Pearson’s correlation test was applied. B. Correlations between changes in STN gamma oscillation and motor disability. Decreased gamma mean power (r = 0.68, *p* = 0.029. Pearson’s correlation test) and peak power (r = 0.74, *p* = 0.015. Pearson’s correlation test) were both positively correlated with decreased success rate. Pearson’s correlation test was applied.

### STN decreased gamma power was positively correlated with the decreased firing rate

ACS caused decreases in STN oscillations and firing rate, prompting us to further investigate potential correlations between these changes. We used Pearson’s correlation test to analyze the correlation between the decreased STN firing rate and the changes in delta and gamma peak power and mean power. The results revealed that a significant positive correlation between decreased STN gamma mean power (r =0.71, *p* = 0.020) and peak power (r =0.78, *p* = 0.007) with the decreased STN firing rate. In contrast, decreased STN delta mean power (r =0.57, *p* = 0.083) and peak power (r =0.31, *p* = 0.388) were not correlated with the decreased STN firing rate (Fig. 8A).

**Fig.8.**
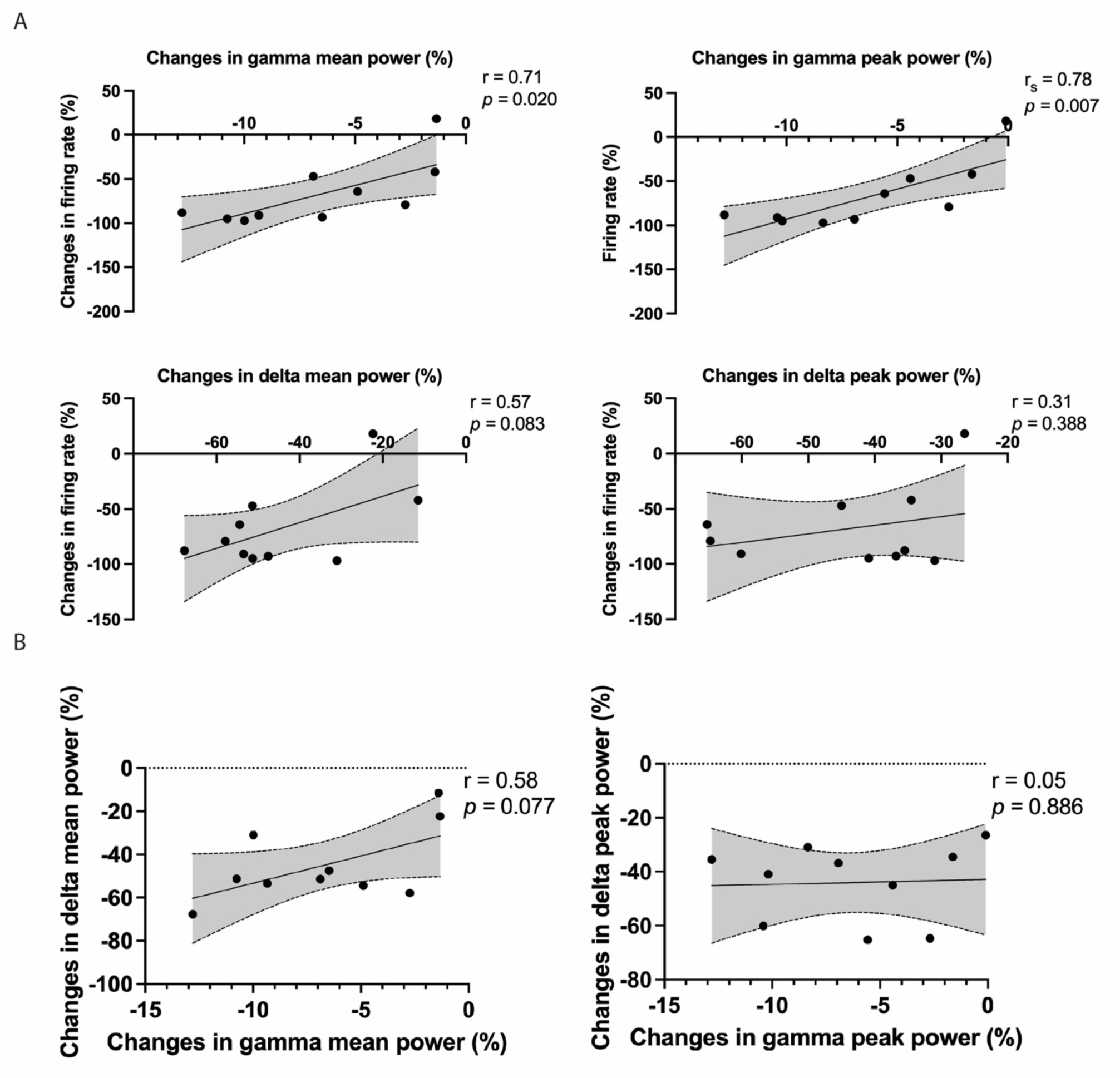
Spearman’s rank correlation results. A. Correlations between changes in STN oscillations and firing rate. Decreased STN gamma mean power (r =0.71, *p* = 0.020) and peak power (r =0.78, *p* = 0.007) are positively correlated with the decrease in STN firing rate. However, decreased STN delta mean power (r =0.57, *p* = 0.083) and peak power (r =0.31, *p* = 0.388) are not correlated with a decrease in STN firing rate. Pearson’s correlation test was applied. B. Correlations between changes in delta oscillation and gamma oscillation. Decreased gamma mean power is not correlated with decreased delta mean power (r = 0.58, *p* = 0.077). Similarly, the decrease in gamma peak power is not correlated with the decrease in delta peak power (r =0.05, *p* = 0.886). Pearson’s correlation test was applied.

### STN decreased delta power was not correlated with the decreased gamma power

ACS caused decreases in STN delta and gamma power. Thus, we questioned whether they were correlated. To address this, we used Pearson’s correlation test to analyze the relationship between the changes in delta peak and mean power and the changes in gamma peak and mean power. The results revealed that decreased gamma mean power was not correlated with decreased delta mean power (r = 0.58, *p* = 0.077). Similarly, the decrease in gamma peak power was not correlated with the decrease in delta peak power (r =0.05, *p* = 0.886) (Fig. 8B).

### Verification of the electrode tip positions

To verify electrode tip positions, we used H&E staining. In Experiment 2, electrodes were successfully implanted in STN of 10 rats in experimental group, with tip positions ranging from 3.12 to 4.08 mm posterior to bregma (Fig. 4I), and in 7 rats in control group, with positions ranging from 3.24 to 3.72 mm posterior to bregma (Fig. 4j). In Experiment 3, electrodes were successfully implanted in 2 rats in experimental group, with tip positions ranging from 3.60 to 3.72 mm posterior to bregma (Fig. 6G), and in 2 rats in the control group, both at 3.72 mm posterior to bregma (Fig. 6H).

## Discussion

In this study, we explored STN neural dynamics following ACS and examined the correlation between changes in STN activity and motor disability. The MC and STN are connected via the HDP, playing a crucial role in controlling motor function. Therefore, we aimed to investigate whether the STN is affected after ACS. To do this we implanted recording electrodes in the rat STN, induced ACS lesion in MC and assessed motor behaviour. Our study is the first to demonstrate that ACS modulates firing rate and oscillations in STN in rats (Fig.1). Notably, this ACS-led STN modulation correlated with motor disability.

Our findings indicate that 1) post-stroke cortico-subthalamic diaschisis (PSCSD) occurs during ACS, resulting in STN inhibition and a decrease in delta (0.5-4Hz) and gamma oscillation (50-140Hz). We refer to this decrease in delta and gamma oscillations as abnormal STN oscillations. 2) Abnormal STN oscillations could serve as a biomarker for motor disability severity as measured by SPRT in acute stage after cortical stroke in rats, as the severity of acute motor disability correlates with abnormal STN oscillation. We will elaborate on these points further below. Based on our findings, we propose the following hypotheses for future research 1) stroke is a neurocircuitry disorder, particularly involving STN and motor cortex. We refer to this neurocircuitry as post-stroke abnormal motor circuit. 2) Motor restoration can be achieved by modulating this abnormal neurocircuitry (e.g., the post-stroke abnormal motor circuit) or modulating the abnormal STN oscillations.

Diaschisis is defined as the distant neurophysiological changes caused by a focal injury.^27^ In our study, we induced a focal ACS lesion in the M1, which causes a significant distant neural inhibition and abnormal oscillations in STN. Thus, we refer to this phenomenon as PSCSD. The formation of PSCSD may occur in two stages: 1) The MC becomes abnormal following ACS and persistently sends abnormal cortical outputs to the STN via the HDP.^12^ 2) The STN receives the cortical abnormal inputs and responds abnormally, culminating in subsequent neurophysiological changes in STN (Fig. 1B). However, further evidence is needed to validate this hypothesis. Furthermore, STN inhibition may occur rapidly after ACS induction based on the observation in Experiment 3. This implies that 1) STN is highly sensitive to ACS injury and responds rapidly to cortical injury during ACS induction, possibly due to the fast-speed intercommunication (6 to 24 m/s) between MC and STN facilitated by HDP.^28^ However, we took this argument cautiously due to the low sample size in Experiment 3. Additional studies are needed to validate this argument. 2) During and after ACS, STN functions abnormally, which may compromise its role in motor control, leading to motor dysfunction and disability. Although we have shown a correlation between motor disability and abnormal STN oscillations, more studies are needed to clarify how this abnormal STN oscillations affect motor control.

Post-stroke STN inhibition may act as a partial compensatory response to ACS. However, it could also lead to pathological consequences. According to a classic model of basal ganglia, the indirect pathway plays a major role in suppressing movement.^29^ In our study, the indirect pathway was inhibited, as evidenced by STN inhibition, suggesting that movement suppression may be reduced. Specifically, the internal globus pallidus (GPi) is expected to be inhibited due to the STN inhibition. This GPi inhibition subsequently leads to thalamic activation, resulting in enhanced activation of some regions of MC that survive the ACS lesion. This aligns with several reports demonstrating that neural activity was found to be elevated focally in the peri-infarct region (5mm from the stroke lesion centre) of MC following stroke.^30, 31^ Thus, we interpret post-stroke STN inhibition as a partial compensatory mechanism, wherein relatively enhanced MC activation compensates, to some extent, for the substantial loss of MC activity caused by ACS. More notably in clinical practice, subthalamic hyper-activation causes bradykinesia,^32^ while STN inhibition mitigates bradykinesia and even causes involuntary movements or hemiballism.^33, 34^ These phenomena were also reported in animals.^35, 36^ In our study, motor disability was evident following ACS. While STN inhibition may facilitate motor initiation,^33–36^ it appears inadequate to fully offset the motor disability observed in the rats of this study. Behaviourally, this suggests that STN inhibition likely represents a partial compensatory response to the motor disability after ACS. However, we have not tested whether reducing or augmenting STN inhibition could exacerbate or improve motor disability following ACS. Therefore, additional evidence is needed. Furthermore, this partial compensatory response may also become pathological in certain cases, turning the peri-infarct region into a zone of hyperactivity and leading to focal post-stroke epilepsy.^37^ In summary, post-stroke STN inhibition may partially compensate for ACS, but it also has the potential to cause pathological effects. Moreover, as STN is part of basal ganglia, it is important to recognise that basal ganglia is highly likely to become dysfunctional due to the cascade effects of STN inhibition (Fig.1B).^29^ Therefore, It is reasonable to shift some focus from MC to basal ganglia.

ACS not only causes STN inhibition, but also causes STN abnormal oscillations. Interestingly, delta and gamma bands have been reported to be associated with motor function.^14–16, 38^ STN gamma oscillation was reported to have prokinetic effects.^39^ Cassidy et al.^16^ noted that in patients with Parkinsonism, STN power is dominated by gamma oscillations, which increase during externally paced movement following dopaminergic stimulation. However, STN gamma activity remains unchanged in the absence of dopaminergic stimulation. This prokinetic effect is further supported by antiparkinsonian (e.g., bradykinesia) benefits observed with high-frequency stimulation of STN, which is believed to increase STN gamma power.^40^ Thus, it is rational that an abnormal decrease of prokinetic STN gamma oscillation may result in motor disability post-stroke. Additionally, Ramanathan et al. indicated that cortical delta power diminished after stroke but showed a recovery during the process of spontaneous motor recovery post-stroke in rodents.^14^ This finding is supported by a clinical study that reported a reduction in delta power in patients with acute stroke, which progressively increased with motor recovery.^15^ Furthermore, cortical gamma power is also linked to motor function.^38^ Thus, we hypothesise that the abnormal decrease in both STN delta and gamma oscillations observed in our study may be driven by or synchronised with decreased cortical delta and gamma power through HDP. This raises the question whether there is an abnormally increased synchronisation in delta and gamma frequency bands between STN and MC that impairs motor function after stroke? If this is the case, the ACS may cause pathological intercommunication through abnormal post-stroke motor circuit between STN and MC, which plays a role in motor disability. Gaining a deeper understanding of this pathological intercommunication and the role of abnormal post-stroke motor circuit in motor disability could provide valuable insights into the condition and inform the development of effective strategies to treat motor disability following stroke. Furthermore, STN may exert detrimental influence on both basal ganglia and MC through the indirect pathway and STN-cortex pathway when it functions abnormally after ACS (Fig.1B).^14, 29^ This STN dysfunction could aggravate motor disability through 1) disrupting motor regulation and control in both basal ganglia and MC, and 2) perpetuating the spread of erroneous signals between MC and basal ganglia, thereby driving flawed neuroplasticity in both MC and basal ganglia, which ultimately aggravates motor disability after ACS.

It remains uncertain whether post-ACS abnormal STN activity causes motor disability. In our study, the abnormal STN activity was observed exclusively in rats with motor disability after ACS induction. Researchers have reported that reducing abnormal beta activity in STN can alleviate motor symptoms in Parkinson’s disease patients.^41^ Thus, it suggests the possibility that abnormal STN activity contributes to motor disability. However, current study has not demonstrated that reducing abnormal STN activity can alleviate motor disability following ACS. Further research is needed to validate this argument. Nonetheless, our findings suggest that is worthwhile to test electrical stimulation of STN for enhancing motor recovery after stroke. This is particularly relevant given that high-frequency electrical stimulation of STN has traditionally been thought to suppress abnormal STN activity, as suggested by the widely accepted #inhibition hypothesis”.^42^

Our results highlight a great potential of using post-stroke abnormal STN oscillations as a biomarker for predicting motor disability in acute stage after cortical stroke in rats. This potential is supported by two observations: 1) abnormal STN oscillations are significantly correlated with motor disability. 2) STN seems sensitive to ACS injury and responds swiftly to it. However, further research is still needed to strengthen this argument, and additional studies are required to explore whether these findings can be translated from bench to bedside.

We found that decreased STN gamma power was significantly correlated with decreased STN firing rate after ACS. This finding is consistent with previous studies indicating decreased gamma power is associated with decreased firing rate.^43^ It has been proposed that gamma power reflects the strength of synchrony of firing activity.^44^ Therefore, it is reasonable that decreased STN firing rate results in a decreased synchrony in gamma band across the STN neural population, as significantly fewer STN neurons are discharging. This decrease in synchrony across the STN neural population ultimately leads to the observed decrease in STN gamma power. Furthermore, both delta power and gamma power were found to be decreased in STN following ACS. This raises a question whether there is a relationship between decreased gamma power and decreased delta power in STN after ACS? Our results revealed no relationship between them, suggesting that these two decreased oscillations may be independent from each other.

There are two main limitations in this study. First, rats were not randomly assigned to control group and experimental group in Experiment 2, potentially introducing bias into the study. The low success rate of electrode implantation into the STN in the pilot study presented challenges in ensuring consistent implantation in Experiment 2. As a result, we first conducted data collection in the experimental group, followed by the control group. However, this success rate improved progressively with increased surgical experience. Thus, we did randomization in Experiment 3. Future studies will aim to implement randomization protocols to enhance the robustness of the study. Second, the sample size is small in Experiment 3, and as such, our results should be replicated in future studies with larger sample sizes to increase the reliability.

In conclusion, our study is the first to show that ACS influences firing rate and oscillations in STN in rats. Interestingly, this ACS-driven modulation of the STN was linked to motor disability. Notably, the abnormal STN oscillations can serve as a biomarker for predicting motor disability during the acute stage following cortical stroke in rats. Furthermore, our findings suggest the potential for developing neuromodulation strategies to alleviate post-stroke motor disability by targeting and reducing abnormal STN activity.

## Acknowledgement

We would like to sincerely thank Prof. Christelle Baunez for her invaluable guidance in implementing the electrode implantation in STN. This work was supported by Fonds Wetenschappelijk Onderzoek’s Strategic Basic Research (SBO) program (S000221N) and the Boston Scientific Chair in Neuromodulation for Stroke.

## Author Contributions

ZD.D. U.K. and B.N. contributed to the conception and design of the study. ZD.D. contributed to the acquisition data. ZD.D., M.M.L., and B.A. contributed to analyze the data. ZD.D., B.A. and B.N contributed to writing the manuscript. ZD. D contributed to preparing the figures. All authors revised this manuscript.

